# Foveolar drusen decrease fixation stability in pre-symptomatic AMD

**DOI:** 10.1101/2023.10.09.560481

**Authors:** J. Murari, J. Gautier, J. Daout, L. Krafft, P. Senée, P. Mecê, K. Grieve, W. Seiple, D. Sheynikhovich, S. Meimon, M. Paques, A. Arleo

## Abstract

**Purpose:** This study aims at linking subtle changes of fixational eye movements (FEM) in controls and in patients with foveal drusen using adaptive optics retinal imaging in order to find anatomo-functional markers for pre-symptomatic Age-related Macular Degeneration (AMD).

**Methods:** We recruited 7 young controls, 4 older controls and 16 presymptomatic AMD patients with foveal drusen from the Silversight Cohort. A high speed research-grade adaptive optics flood illumination ophthalmoscope (AO-FIO) was used for monocular retinal tracking of fixational eye movements. The system allows for sub-arcminute resolution, high-speed and distortion-free imaging of the foveal area. Foveal drusen position and size were documented using gaze-dependent imaging on a clinical-grade AO-FIO.

**Results:** FEM were measured with high precision (RMS-S2S=0.0015° on human eyes) and small foveal drusen (median=60µm) were detected with high contrast imaging. Microsaccade amplitude, drift diffusion coefficient and ISOline Area (ISOA) were significantly larger for patients with foveal drusen compared with controls. Among the drusen participants, microsaccade amplitude was correlated to drusen eccentricity from the center of the fovea.

**Conclusions:** A novel high-speed high-precision retinal tracking technique allowed for the characterization of FEM at the microscopic level. Foveal drusen altered fixation stability, resulting in compensatory FEM changes. Particularly, drusen at the foveolar level seemed to have a stronger impact on microsaccade amplitudes and ISOA. The unexpected anatomo-functional link between small foveal drusen and fixation stability opens up a new perspective of detecting oculomotor signatures of eye diseases at the presymptomatic stage.

## Introduction

Age-related macular degeneration (AMD) is the main cause of vision loss in developed countries, affecting 180M people in 2020 with an expected prevalence of 288M by 2040^1^. AMD starts with the accumulation of extracellular deposits, called drusen, in-between the Bruch’s membrane and the retinal pigment epithelium (RPE). Multiplication and expansion of drusen is associated with late stages of AMD, which are classified into exudative (wet-AMD) and/or atrophic (dry-AMD)^2^. Wet-AMD is caused by irregular blood vessel growth (i.e., neovascularization) at the macular level, and it affects approximately 10-20% of AMD patients^3^. Although macular neovascularization can be successfully treated using anti-VEGF therapies, these treatments do not prevent the development of atrophy, which causes sight loss in the long term^4^. Geographic atrophy or dry-AMD is the irreversible loss of photoreceptors caused either by drusen development or neovascular growth, and it accounts for about 80 to 90% of the cases^2^. No effective treatment currently exists for this form of AMD. The absence of efficient treatment strategies for the late-stage AMD is exacerbated by the fact that patients frequently remain unaware of their AMD condition^5^, so that they typically receive a diagnosis only upon the emergence of clinical indications, often at a stage too advanced for effective implementation of preventive strategies. There is therefore an urgent need to characterize anatomo-functional relationships associated with the earliest asymptomatic AMD stages aiming at complementing existing medical imaging techniques with functional measures of retinal anomalies that can serve as biomarkers of future disease onset. If indeed the pre-symptomatic structural anomalies are associated with functional, e.g. oculomotor, changes, their timely detection may also incite the individuals with a risk of developing AMD to engage in risk mitigation strategies to delay AMD onset and progression^6,7,8^.

The objective of the present study is to test the novel hypothesis that the appearance of drusen, an early structural anomaly in presymptomatic AMD, is associated with detectable changes in microscopic eye movements during ocular fixation. Even though the development of early-to-late stages of confirmed AMD has already been associated with changes in fixation stability, decreased visual acuity, reduced visual field, slower reading speed, lower contrast sensitivity, and an overall deterioration of functional vision useful for daily activities^9–11^, these previous studies did not explore fixational eye movements in presymptomatic stages, nor did they characterize the link between retinal structure changes and fixational eye movements. In patients with central scotomas, fixation stability as assessed by retinal motion compensation strategies has been shown to worsen with AMD progression^12^. Moreover, advanced geographic atrophies were reported to affect different components of fixational eye movements, such as microsaccades (small saccades of amplitude below 1°) and ocular drift^13,14^ (random Brownian-walk-like eye movements between microsaccades). More specifically, these studies report higher microsaccade amplitudes associated with macular lesions^13^ and lower spectral whitening of natural scenes by ocular drift in patients with long-lasting macular disease, compared to age-matched controls^14^. Given the significant changes in fixational stability in advanced forms of AMD, it is important to test whether even the first signs of potential future disease can induce changes in eye movement statistics.

The hypothesis was tested by simultaneously performing *(i)* high-resolution retina imaging to visualize and characterize small asymptomatic alterations of the foveal structure, *(ii)* high-speed retina tracking to detect fixational eye movements. We studied the relationship between subtle structural retinal anomalies and fixational eye movement characteristics by examining healthy controls and participants with foveal drusen, but no scotoma or atrophy according to conventional measures. If the appearance of first drusen indeed leads to fixation instability, as the current study shows, such a structure-function relationship provides the possibility to use fine oculomotor measurements to predict the appearance of retinal anomalies and therefore contribute to the development of early anatomo- functional biomarkers for urgently needed screening approaches in presymptomatic AMD.

## Methods

### Participants

A cohort of 39 participants were recruited for the study as part of one of three groups: young controls (n=7, [2 females], age=31±7 yo), older controls (n=20, [14 females], age=75±4 yo) and drusen participants (n=12, [8 females], age=79±3 yo). The initial assignment of participants to the groups relied on the detection of drusen by standard spectral-domain optical coherence tomography (SD-OCT) imaging (Spectralis OCT, Heidelberg Engineering, Germany). All participants were recruited from the SilverSight cohort^15^, and they were fully screened and ascertained not to suffer from any age-related sensory, cognitive, or motor pathology.

Prior to the beginning of the experiment, older controls and drusen participants were screened using a novel imaging method, developed previously in our laboratory using Adaptive Optics Flood Illumination Ophthalmoscope (AO-FIO, rtx1, Imagine Eyes), which is capable of detecting foveal drusen not visible in standard SD-OCT or fundus imaging^16^ (see the next section). As a consequence, 9 participants that had initially been recruited as older controls were actually found to have drusen in the foveal region, and they were thus moved into the drusen group. In addition, a total of 12 participants had to be excluded from study: 3 because of too blurry retina images on the rtx1; 3 due to poor signal-to-noise ratio during the fixation task; 2 because they were not fixating the right target; 3 because of excessive tiredness and discomfort during the experiment; and 1 because she exhibited fixation patterns typical of another disease. As a result, a total of 27 participants were kept for the analyses: 7 young controls ([2 females], age=31±7 yo), 4 older controls ([1 female], age=72±2 yo), and 16 drusen patients ([10 females], age=78±4 yo). Their demographic and ophthalmological characteristics are given in **Table 1**.

| Tag | Age (yo) | Tested eye | Visual acuity | Number of drusen | Mean drusen area ± std (arcmin<sup>2</sup>) | Total drusen area (arcmin<sup>2</sup>) | Mean drusen eccentricity ± std (arcmin) |
| --- | --- | --- | --- | --- | --- | --- | --- |
| DI1 | 79 | OD | 1.2 | 7 | 79 ± 14 | 556 | 27 ± 22 |
| DI2 | 78 | OD | 1 | 7 | 169 ± 66 | 1182 | 28 ± 23 |
| DI3 | 80 | OD | 1 | 5 | 75 ± 18 | 376 | 30 ± 25 |
| DI4 | 84 | OS | 1.2 | 7 | 143 ± 32 | 998 | 37 ± 29 |
| DI5 | 80 | OD | 1.2 | 32 | 339 ± 147 | 10861 | 42 ± 31 |
| DI6 | 83 | OS | 1 | 12 | 176 ± 50 | 2121 | 42 ± 26 |
| DI7 | 76 | OS | 1 | 9 | 131 ± 72 | 1184 | 45 ± 27 |
| DI8 | 74 | OD | 1 | 2 | 106 ± 29 | 212 | 45 ± 14 |
| DI9 | 75 | OD | 1.2 | 6 | 178 ± 40 | 1069 | 56 ± 32 |
| DO1 | 73 | OD | 0.8 | 5 | 148 ± 78 | 741 | 62 ± 22 |
| DO2 | 84 | OS | 1 | 13 | 177 ± 48 | 2313 | 63 ± 25 |
| DO3 | 70 | OS | 1 | 15 | 180 ± 70 | 2710 | 62 ± 32 |
| DO4 | 80 | OD | 0.7 | 6 | 128 ± 49 | 766 | 68 ± 19 |
| DO5 | 81 | OD | 1.2 | 20 | 139 ± 39 | 2791 | 69 ± 42 |
| DO6 | 74 | OS | 0.8 | 14 | 59 ± 15 | 826 | 76 ± 25 |
| DO7 | 76 | OD | 1.2 | 28 | 189 ± 75 | 5311 | 92 ± 34 |
| Mean for drusen group (n=16) | **78 ± 4** | **/** | **1 ± 0.2** | **11 ± 8** | **133 ± 59** | **2126 ± 2000** | **53 ± 27** |
| Mean for older controls (n=4) | **72 ± 2** | **/** | **>1** | **/** | **/** | **/** | **/** |
| Mean for young controls (n=7) | **31 ± 7** | **/** | **>1** | **/** | **/** | **/** | **/** |

All clinical, experiment, and data management procedures were approved by the Ethical Committee “CPP Ile de France V” (ID RCB 2015-A01094-45, n. CPP: 16122 MSB), and they adhered to the tenets of the Declaration of Helsinki. A signed consent was obtained by each participant to be included in the study as well as for the inclusion in the Multimodal Ophthalmologic Imaging protocol (IMA-MODE, ID\_RCB 2019-A00942-55 CPP: Sud Est III 2019-021-B) of the Quinze-Vingts National Ophthalmology Hospital.

### Drusen detection by gaze-dependent imaging

In order to reliably detect the presence of even the smallest drusen, gaze-dependent fundus imaging of all older participants’ central retina was performed using a commercial clinical-grade AO-FIO (rtx1, Imagine Eyes) following the method and protocol described by Rossi et al.^16^. Compared to direct single-shot retinal flood imaging, gaze dependent imaging computes the local contrast maxima across a stack of 5 images whose gaze direction has been sequentially shifted: one 4°x4° fovea-centered image of the retina, focused on the photoreceptors, and four 4 horizontally and vertically displaced images (±1° with respect to the center image). After image registration, the maximum of the standard deviation for each x, y pixel of the 5-image stack is computed to make gaze-varying structure visible. In the resulting contrast image, drusen and their delineation are clearly visible (**Fig. 2c**). This gaze-dependent retina imaging method allows for the detection of foveal drusen less than 25 μm in diameter that are not detected by standard imaging methods and it also visualizes much more reliably drusen 25-60 μm in diameter, providing more precise information about their size and position^16^. In a final step, drusen were manually annotated as ellipses by two experts in medical images, followed by a validation by two ophthalmologists (including M.P.). All the image filtering, registration, and processing were performed using custom software written in MATLAB (Mathworks, USA, 2023).

### Ocular fixation task

Following drusen assessment, all participants were tested in an ocular fixation experiment in order to see whether the presence of drusen affects fixational eye movements. In order to precisely detect even the smallest eye movements, video recordings of the fovea during fixation were performed using the PARIS AO-FIO (ONERA)^17^ located at the Quinze-Vingts National Ophthalmology Hospital in Paris, France. Both the PARIS AO-FIO retinal imaging setup and retinal motion tracking process are described in the following sections. Before the ocular fixation task, the dominant eye was dilated with a drop of 1% tropicamide (*Mydriaticum*, tropicamide 0.5%). The recordings were performed in a dark room with minimal stray light and with the participant’s head fixed on a chinrest. The fixation task consisted of two phases. In the initial phase, participants were required to fixate a central cross for 5 seconds. This phase served to initialize the retinal motion tracking procedure. In the experimental phase, the participants were asked to fixate a circle (outer diameter: 17 arcmin (0.28°), inner diameter: 11 arcmin (0.18°)) for 30 seconds, across 5 trials. The instructions were *“Please, fixate the circle for 30 seconds. Do not speak and try not moving your head. Do not hesitate to blink if you need to.”,* and a vocal indication about time progression was provided after 15 seconds.

### Retina imaging setup

High-resolution structural imaging of the fovea was performed using the PARIS AO-FIO^17^, through video recordings focused on the photoreceptor layer. Concurrently with retina imaging, the Paris AO-FIO allowed for the projection of NIR illumination patterns (i.e., stimuli) onto the retina via a deformable Digital Micromirror Device (DMD)^18^ as well as for partial-field illumination^19^. The stimulus projection with the DMD was controlled using a custom-made Python code and the ALP4Lib python library wrapper. The PARIS AO-FIO system consists of two optical channels: the Wavefront correction channel, and the illumination and imaging channel, combined with the stimulation channel. A fibered SuperLuminescent Diode (Exalos, Switzerland) with nominal center wavelength of 750 nm measures optical aberrations, and an 850nm LED source (Thorlabs, USA) coupled with a liquid fiber homogeneously illuminate the retina over a 4° diameter field of view (**Fig. 1a**). The adaptive optics compensated for the spherical, cylindrical and some higher order aberrations of the eye’s optics before the stimulus reached the retina. The backscattered light from the eye was reflected back on the deformable mirror before reaching the imaging camera. The adaptive optics loop was controlled with a LabVIEW program and image acquisition was monitored via the Holovibes software^20^.

**Figure 1.**
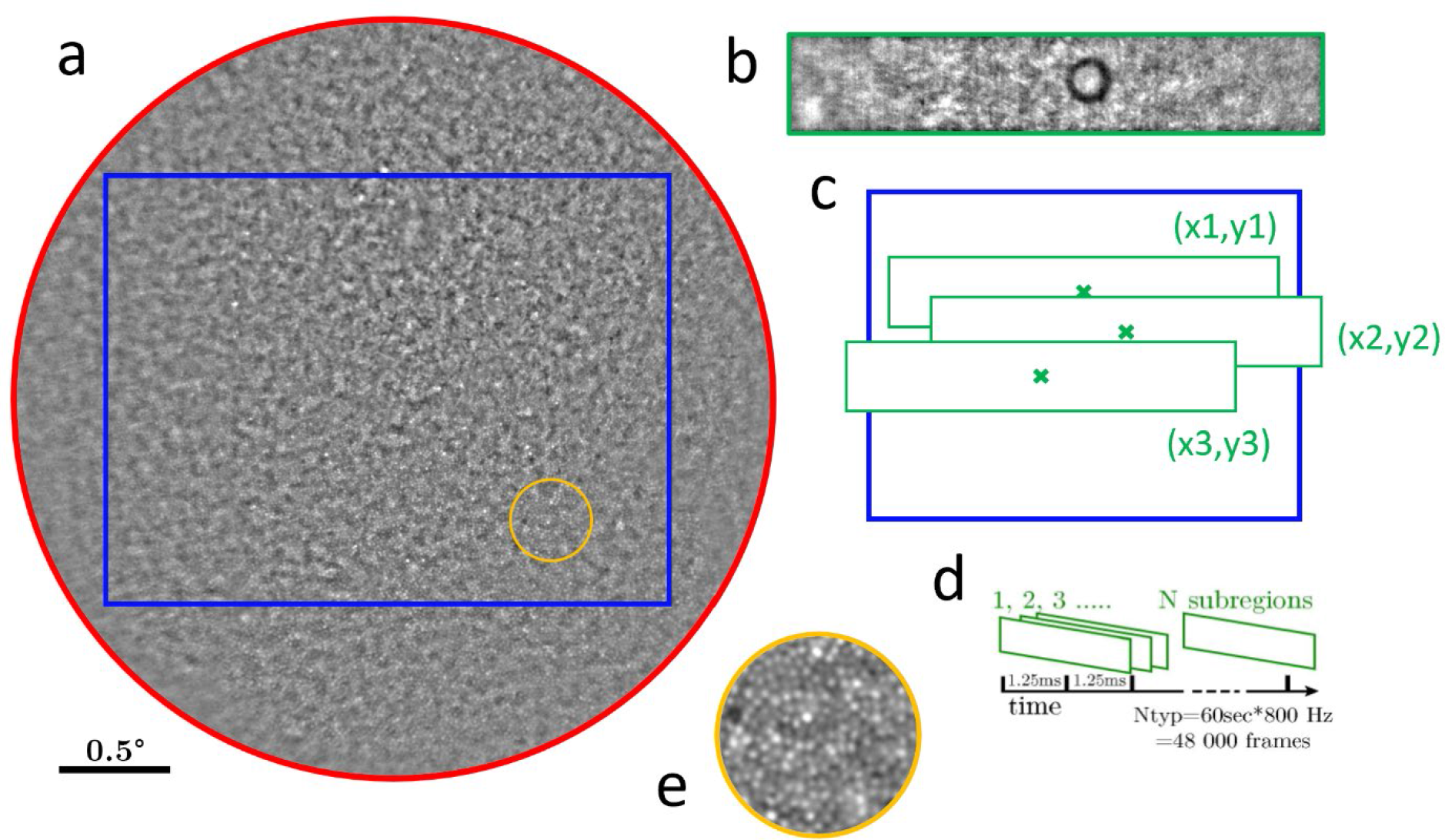
Principle of the AOFIO retinal tracking. (a) The 4° full field obtained after initial phase of the fixation task (5 sec at 100 Hz, in red). Only the rectangular sub-part (in blue) was used as a reference image for subsequent eye movement analysis, as it is enough to track and monitor eye movements during central fixation (typically less than 2° in amplitude). During the experimental phase of the task, the 2 x 0.5° subregion (in green) is tracked at very high speed (800 Hz, 16 bits). (b) An actual frame recorded at this frequency. (c) Eye displacement is obtained by phase-correlation over the reference image. (d) Custom eye tracking software processes large sequences of up to 5 Mbits/frame. (e) A 3x zoom on an area 1° away from the fovea with clearly visible photoreceptor mosaic.

**Figure 2.**
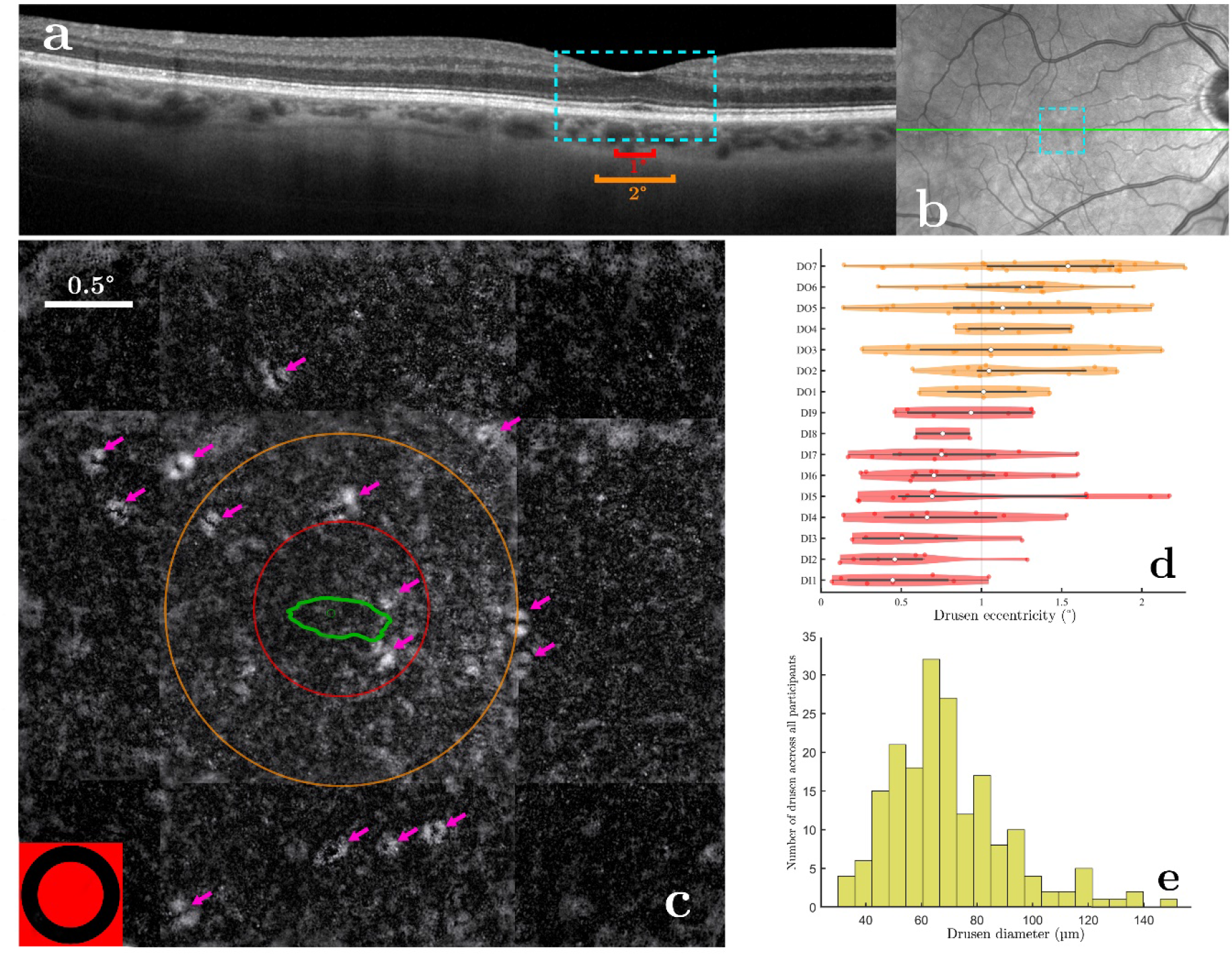
Drusen visualization and identification. (a) SD-OCT image of a patient. A 2x zoom on the 4° center area is shown in blue. The red and orange lines are the same size as the circle diameters in (c). (b) Fundus image with the OCT cut of (a) in green and the blue dotted area showing the 4° zone where gaze-dependent imaging is performed. (c) Gaze-dependent image of the 4°x4° zone around the fovea of the same patient. Pink arrows indicate the drusen: the white ring is the slope of bump cause by the druse the while the black center part is the tip. The red and orange circles are respectively of 0.5° and 1° of radius. The green line is the ISOA 86% On the bottom left corner the fixation target to scale. (d-e) Distribution of drusen eccentricities (c) and diameters (d) across all participants.

### Retina motion tracking

During the initial phase of the ocular fixation task (5s), a full field sequence of images covering a 4° diameter field of participants’ retina was recorded at 100 Hz (2048 x 2048 pixel frames). The recorded full field retinal images were first stabilized and then averaged into a unique squared full field reference image of the center of the retina (**Fig. 1a**). Then, during the experimental phase of the task, only a sub-band (2° x 0.5°, 1400 x 256 pixel, with the black circle target at the center) was recorded at 800 Hz to be used for retina motion tracking (**Fig. 1b**). Retinal motion was tracked using a phase-correlation registration method^21,22^ (**Fig. 1c**) after bandpass filtering the image sequences. The algorithm consisted in finding the position that best fitted each frame of the high-speed video onto the reference image. The best fit was considered as the peak value coordinates of the correlation map computed in the Fourier domain. See **Clip 1**.

Even if the maximum sampling frequency of the camera was only limited by vertical pixel resolution, we found that 1400-pixel horizontally offered a good tradeoff between field coverage and video bandwidth. The resulting 0.684 MB/frame (16 bits) led to a very large 547 MB/s video bandwidth that could only be recorded with solid state hard drives and ad-hoc recording software (Holovibes). Videos were recorded on a separate computer than the one used for stimulus presentation.

### Detection of retina motion related events

Processing of the raw pixel displacement of each frame relative to the reference images to ocular movement in arcminutes with annotated events was made with a MATLAB custom program (based on the work of Bowers and Gautier^23^). Microsaccades were detected using an acceleration threshold inspired by Engbert^24^. An acceleration, rather than velocity, threshold (factor N=3 standard deviation of the median) was used in order to differentiate fast ocular drift epochs from small and slower microsaccades which can have similar velocities but distinct accelerations. The acceleration was computed for the 30 s fixation, and smoothed using running average with a 70∼ms window. This adaptive algorithm allowed for thresholds that fitted the participants who had relatively small microsaccades and stable ocular drift, as well as the participants with larger microsaccade and drift amplitudes. In addition, to ensure that microsaccades were all entirely detected and that some parts of ocular drift were not mislabeled as microsaccades, all traces were manually verified after the automatic annotation. Blinks were detected automatically as sudden drops in mean frame pixel intensity, which corresponded to the closed eye reflecting less light back to the camera. Rejected data were epochs of ocular coordinates marked by the user as false data due to failures in the imaging or the tracking (to the exclusion of blinks). They accounted for 0-3% of the fixation data for each participant. Everything in-between the three types of events – microsaccades, blinks or rejected data – was automatically marked as drift.

Root mean square sample to sample (RMS-S2S) deviation was calculated to measure the precision of the retina tracking method. A given RMS-S2S of an eye-tracker quantifies the average movement due to noise between successive data is its squared value^25,26^. Filtering ocular traces can have a strong effect on RMS-S2S and skews the results^27^. However, in this tracking method, ocular traces were used raw, and RMS-S2S appeared to be the right metric to measure precision.

### Estimation of fixation stability

The Bivariate Contour Ellipse Area (BCEA) and, more recently, the ISOline Area (ISOA) metrics have been used to clinically measure fixation stability of patients with central vision loss, including AMD^28^. ISOA is more robust than BCEA as it can delineate non-ellipsoidal fixation patterns^29^ and it can be used to detect possible multiple retinal fixation loci (PRLs), such as in AMD patients. The choice of the bandwidth and the extent (i.e., number of standard deviations) to compute the ISOA is relatively arbitrary and open to criticism^29–31^. The choice of a 1° window for kernel density estimation, which subtends ISOA calculation, has been shown to be robust and appropriate with respect to the classical definition of the 1° foveola^30,31^. Concerning the ISOA contour extent, we used both the standard 68% ISOA (i.e., encompassing 68% of the data points) and the 86% ISOA (i.e., computed over 2 standard deviations in the 2D space defined from the center of the fixation distribution^32^. The rationale was that the 86% ISOA contour would provide complementary insights on the global fixation pattern that might better differentiate healthy controls from participants with foveal drusen.

### Drift diffusion coefficient and amplitude spectrum computation

In the framework of Brownian motion^33,34^, the eye movement during drift epochs can be modeled by a random walk, and motion statistics can be measured by the diffusion coefficient *D*. We computed the diffusion coefficient *D* by least square best fit to the measurements of the empirical standard deviation of the eye displacement as a function of time^35^. The motion amplitude spectra of the ocular drift segments were analyzed by using a multitaper spectral analysis. This analysis was carried separately on the horizontal and vertical components of eye motion^36,37^, on overlapping segments of 1 s. The time half-bandwidth product was fixed at 2.5, and the last taper was dropped to maximize spectral concentration ratios in the Slepian sequences.

### Statistical analysis

Linear mixed-effect model fit by maximum likelihood (*fitlme* function on MATLAB) was used to compare FEM components between groups with the formula (1). *FEM_component_*were microsaccade amplitudes, ISOAs or drift coefficients.

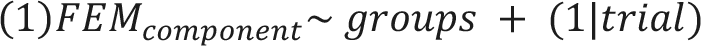

## Results

### Detection and characterization of foveal drusen

Figure 2 shows foveal drusen detected in one participant by using gaze-dependent retinal fundus imaging protocol^16^ (Fig 2c), as well as the corresponding region in an SD-OCT cut across the fovea (Fig 2a**-b**). Overall, in 9 out of 20 participants (45%), initially deemed to have no drusen and therefore assigned to the control group based on SD-OCT images, the presence of foveal drusen was detected using the gaze-dependent detection procedure. These participants were therefore reassigned to the drusen group (see Methods). Across all participants of the drusen group, we quantified the number of foveal drusen, their eccentricity with respect to the center of the fovea, their areas, their diameters (i.e., computed as the average diameter of the ellipses), and the total area covered by all drusen (**Table 1**). The median drusen diameter in the study was 60 µm (0.2°) (Fig 2e).

### Precision and robustness of retinal motion tracking

Retinal tracking during ocular fixation was performed using the Paris AO-FIO^17^ as described in Methods. Figure 3 shows the example traces recorded during 4s of the ocular fixation task in one control participant and one participant with ocular drusen. Our retinal tracking method allowed both microsaccades (shown in magenta) and ocular drift (shown in blue) to be easily visualized and detected in both control and drusen participants.

**Figure 3.**
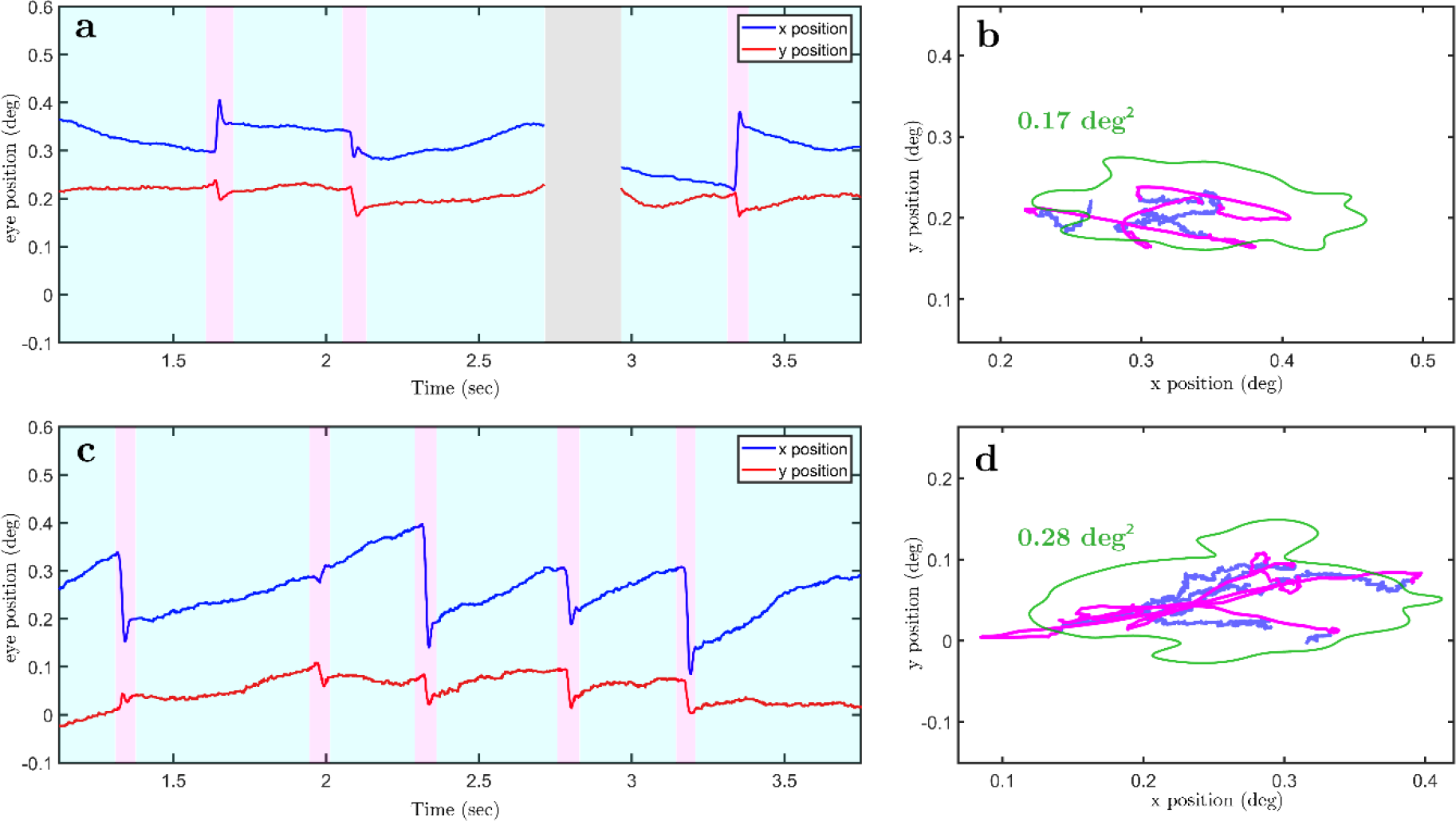
Retina motion traces of a healthy young subject (a-b) vs a subject with drusen (c-d) with the corresponding movement in the visual field. The eye traces are shown for 3 seconds. Microsaccades are annotated in pink and drift in light blue. Eye position is shown in degrees. The green line is the ISOA 86%.

The precision of the tracking procedure was quantified by the average RMS-S2S and achieved 0.0015°, as measured on unfiltered data across all participants. This value is an order of magnitude lower than what is deemed sufficient to detect microsaccades^25^ (0.03°). A low standard deviation of the tracking error (σ=0.0006°) suggests that the method is very robust from participant to participant. Finally, in order to assess possible noise sources stemming from either mechanical vibration of the Paris AO-FIO setup or defects of the tracking algorithm, as well as to get a standard RMS-S2S value, we quantified the precision of the tracking method on non-moving artificial eye (OEMI-7 Ocular Imaging Eye Model). We found an RMS-S2S of 0.00019° (σ=0.00009° over 4 recordings). The precision found on human eyes and the artificial eye is equal or better than any reported values in literature (0.0069° on a DPI Gen5.5 with an non-moving artificial eye^38^ and 0.012° on a human eye^39^; 0.0012-0.0045 on the EWET1 prototypes on human eyes^39^) without being affected by lens-wobble artifacts such as in a DPI or distortion artifacts such as in an AO-SLO system (no RMS-S2S values found for AO-SLO in literature).

The sensitivity to noise was additionally assessed by power spectrum analysis. Power spectra were only weakly affected by noise across all subject groups, even at higher frequencies (above 100Hz, **Figs 5c,d**). The power spectrum of the still model eye was one to two orders of magnitude below human participants (**Supp.** Fig. 1), suggesting negligible setup-related noise.

An addition to be very precise, the method was also found to be resistant to variations in lens transparency (**Supp.** Fig. 2). More specifically, in 11 participants (41%), classified by orthoptists as having corticonuclear lens opacities, the mean RMS-S2S was 0.0018°, only slightly superior to the mean RMS-S2S of all other participants (0.0014°).

### Effect of eccentricity of foveal drusen on microsaccades

To see whether the presence of foveal drusen is associated with changes in eye movements, we plotted individual distributions of microsaccade amplitudes sorted from lowest to highest mean amplitude (Fig 4a). The distributions of young and older participants from the control group were shifted toward the lowest amplitudes, whereas those from the drusen group towards the highest ones, suggesting a link between drusen properties and FEM characteristics. Strikingly, we also noted that the highest microsaccade amplitudes were associated with the presence of drusen mainly inside the foveola (the 0.4-0.6° foveal pit)^40^, suggesting that drusen eccentricities may play an important role in oculomotor changes induced by the presence of drusen.

**Figure 4.**
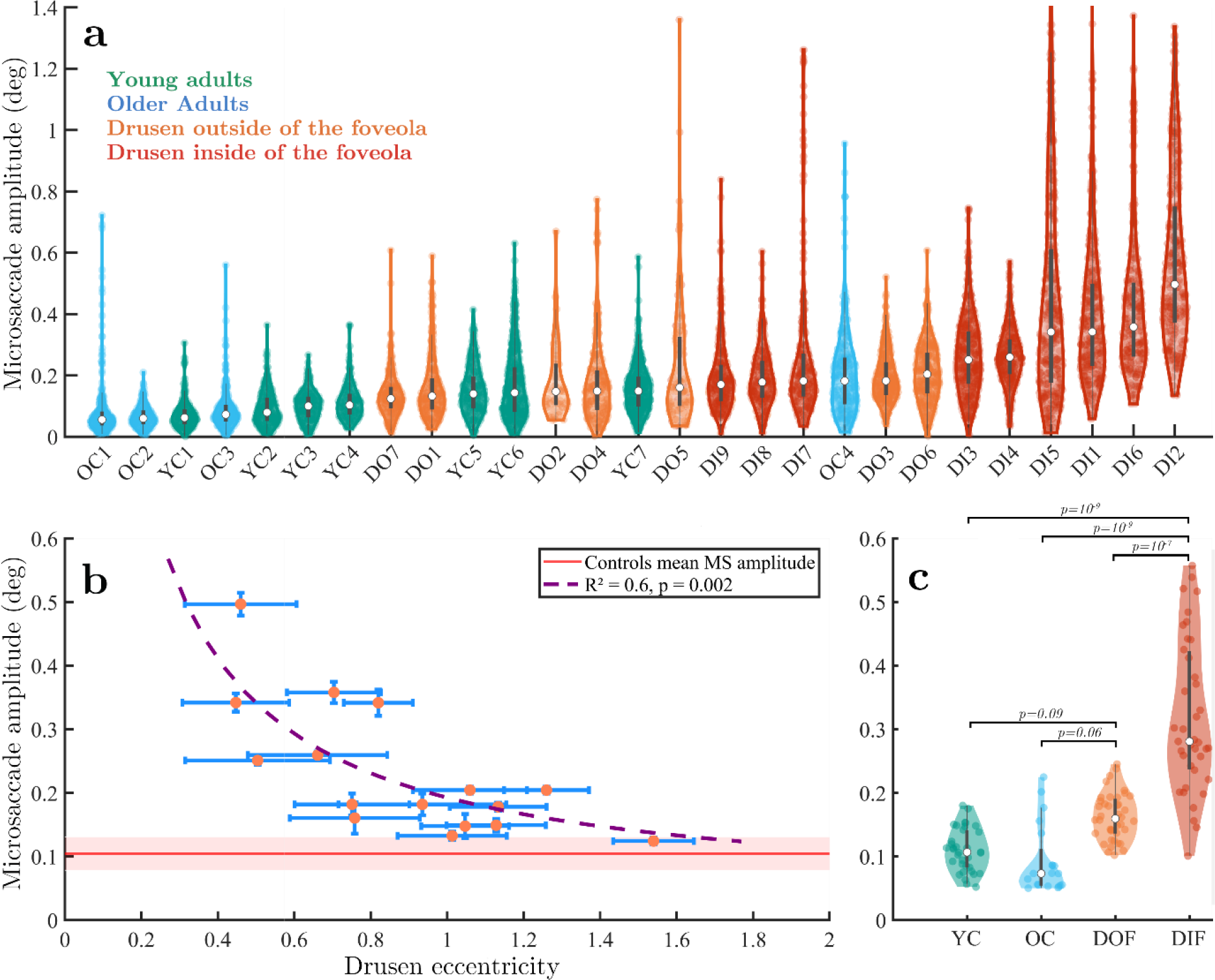
**Microsaccade characteristics across individuals and groups**. **(a)** Violin plots of individual distributions of microsaccade amplitudes across all participants, color-coded by group and sorted with respect to the mean microsaccade amplitudes. Each dot is one microsaccade during the fixation task. (b) Microsaccade mean amplitude with respect to drusen eccentricity from the center of the fovea. Standard errors are shown in blue bars. (c) Microsaccade amplitudes across all four groups. Each dot is the average microsaccade amplitude for 30 seconds (one trial).

**Figure 5.**
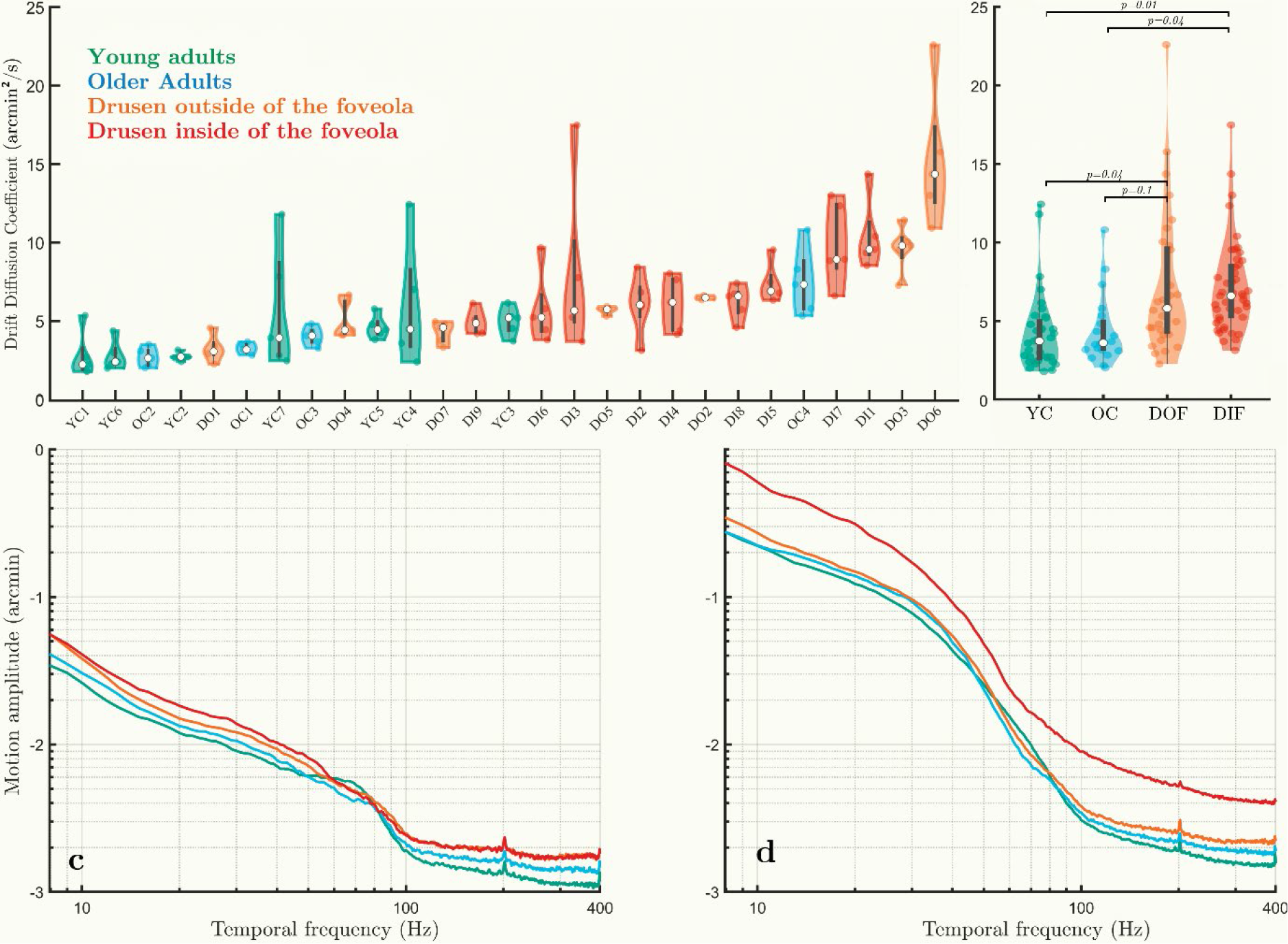
Drift dynamics and motion spectra across groups – horizontal motion. (a) Drift diffusion coefficient distribution across all participants, sorted from lowest to highest. Each dot is one trial. (b) Drift diffusion coefficient across groups (c) Motion spectrum of drifts only across the four groups. (d) Whole motion spectrum (drifts + microsaccades) across the four groups. An increase in dynamic across most frequencies is visible with age, and even further with the presence of foveal or foveolar drusen. This progressive increase is particularly visible in the Mid-frequency (10-40 Hz, top left inset) and very-high frequency range (100-400 Hz, top-right inset).

To quantify this effect, we computed Pearson’s correlation coefficients between the measured drusen characteristics (i.e. their number, area and eccentricity) and microsaccade sizes. We found a significant negative correlation between the participants’ drusen eccentricities and their microsaccade amplitudes (r=-0.74, p=0.001). Hence, the closer drusen were to the center of the fovea, the larger were microsaccades. Conversely, the more eccentric were the drusen in a particular participant, the more similar this participant was to the control subjects in terms of the microsaccade amplitude. No significant correlation was found between the number of drusen, the mean drusen area or the total drusen area and microsaccade amplitude (r=-0.003, p=0.99; r=0.16, p=0.53; r=0.13, p=0.61, respectively. The results in Fig. 4b show that microsaccade amplitudes start increasing exponentially when drusen are located closer than 0.8°-1° with respect to the center of the fovea.

To further characterize the role of drusen eccentricity, participants from the drusen group were subsequently classified into two subgroups. The classification took into account the approximate diameter of the foveola (0.4-0.6°^40^), of the stimulus (0.28°), and of the median foveal druse (0.2°). The first subgroup (named DIF, for “drusen inside the foveola”) included the participants with drusen mainly located in the foveola (i.e., mean drusen eccentricity less than 1° with respect to the center of the fovea, n=8). The other subgroup (named DOF, for “drusen outside the foveola”) consisted of participants with drusen located mainly outside their foveola (n=8).

Statistical analysis using linear mixed effects model revealed significant differences between the microsaccade amplitude in the DIF group and the three other groups (Fig. 4c; DIF vs young controls: |t|=6.5, p=10^-9^; DIF vs older controls: |t|=6, p=10^-9^; DIF vs DOF: |t|=5, p=10^-7^). The group with drusen outside the foveola (DOF) tended to have larger microsaccades than young controls (|t|=1.7, p=0.09) and older controls (|t|=1.8 p=0.06). There was no difference between young and older controls (|t|=0.4, p=0.66).

While no significant differences were found between groups in terms of *average* drusen area, we hypothesized that large drusen closer to the center of the fovea should have an even stronger effect on microsaccade amplitude. The rapid drop-off of both cone density and acuity in the fovea suggest that both drusen area and eccentricity can have tremendous effect on foveal vision. We therefore computed the correlation between drusen area divided by eccentricity (that we called drusen density) and measured FEM characteristics. This new measure was indeed stronger correlated with microsaccade amplitude (r=0.82, p=0.0001) than the eccentricity alone. There were no statistical differences in the rate of microsaccades between groups and all participants produced approximately 1-3 microsaccades per second.

The above results show that drusen eccentricity, a simple property of retinal structure associated with presymptomatic stage of AMD, is a crucial factor explaining the difference in microsaccade size between control participants and subjects with a relatively elevated risk of developing the disease. The quality of this simple structural measure in explaining functional changes can be further improved by taking into account drusen area.

### Impact of foveal drusen on ocular drift and eye motion spectrum

Apart from microsaccades, the ocular drift component of FEM (see Fig. 3) was also affected by the presence of foveal drusen, but to a lesser extent. Indeed, the analysis of ocular drift in the four participant groups defined in the previous section revealed significant differences in drift diffusion coefficients between participants with drusen in the foveola and control subjects (**Figs. 5a,b**; DIF vs young controls: |t|=2.5, p=0.01; DIF vs older controls: |t|=2, p=0.04). Statistical pairwise comparisons between all other groups have shown that the DOF group was significantly different from young controls (|t|=2, p=0.04), but not from older controls (|t|=1.7, p=0.1). No differences were found between young and older controls (|t|=0.1, p=0.93, the absence of statistical differences here should be taken with caution considering the low sample size of older controls). Individual diffusion coefficient revealed larger variability within groups but less between groups (Fig 5a**,b**) when compared to mean microsaccade amplitude (Fig 4a). No significant correlations were found between drift diffusion coefficient and drusen properties across all subjects (eccentricity: r=0.09, p=0.72; drusen number: r=-0.003, p=0.99; mean drusen area: r=-0.1, p=0.69; total drusen area: r=-0.05, p=0.85) A different way to analyze FEM changes induced by drusen presence is look at the motion spectrum across the subject groups. The spectral analysis of ocular drift revealed a progressive increase in motion amplitude both with age and drusen centrality (Fig. 5c), marked in the middle frequency range (10-40 Hz). No difference in motion amplitude was observed at high frequencies (100- 400 Hz) between the two drusen groups, in contrast to an age-related difference in control subjects (Fig. 5c). When accounting for the spectral changes caused by microsaccades (Fig. 5d), the foveolar drusen group clearly stood out across all motion frequencies, potentially impeding later-stage neural processing of the visual input^35^. The differences between the two motion spectra (Fig. 5c**,d**) reflect the statistical differences in drift diffusion coefficient between subjects groups presented above and highlight the more important role of microsaccades, compared to ocular drift, in FEM changes induced by the presence of central drusen.

### Foveolar drusen impair fixation stability

Fixational movements of the eye, including microsaccades and drift, are the primary determinants of fixational stability, a functional measure of ocular stability at the macroscopic level usually evaluated by such measures as ISOline Area^28,29^ (see Methods). The discovered relationship between central drusen and microsaccade size, described in the previous section, suggests that a similar relationship should exist between drusen eccentricity and ocular stability in the fixation task. Indeed, we found a strong and significant negative correlation between ISOA (at 86% level) and drusen eccentricity (r= -0.72, p=0.001) as well as highly significant positive correlation between ISOA and drusen density (r= 0.72, p=0.0004 across all subjects. We checked that there were no significant differences between participants in terms of microsaccade frequencies that could explain changes in fixation stability. In order to better understand the effect of drusen on fixation stability and its topological characteristics, we quantitatively compared average ISOA values across the four groups of subjects using linear mixed modeling approach and visualized average ISOA contours. The participants with drusen mainly in the foveola had the largest ISOA as compared to the other three groups (DIF vs young controls: |t|=5.5, p=10^-7^; DIF vs older controls: |t|=4.9, p=10^-6^; DIF vs DOF: |t|=5.4, p=10^-7^). ISOAs remained similar between the DOF group and young controls (|t|=0.24, p=0.81) or older controls (|t|=0.24, p=0.67), and between young and older controls ((|t|=0.2, p=0.83). These results extend our conclusion from the previous section by showing that impaired fixation stability in subjects with drusen in foveola is primarily mediated by the increase in microsaccade size and not by other FEM properties, such as ocular drift.

Given the consistent and strong effect of foveal drusen on both fixation stability and microsaccade amplitude, it is not surprising that there was a strong correlation between the ISOA 86% values and microsaccade amplitudes across the entire study population (Fig. 7a; r²=0.85, p=0.001). Interestingly when looking at figure 6, in all groups except DIF, higher values for ISOA seemed to come with both proportional increases in microsaccade amplitude and drift diffusion coefficient at the same time. However, for the DIF group, drift diffusion coefficient appeared to lose its correlation with ISOA compared to the rest of the study population.

**Figure 6.**
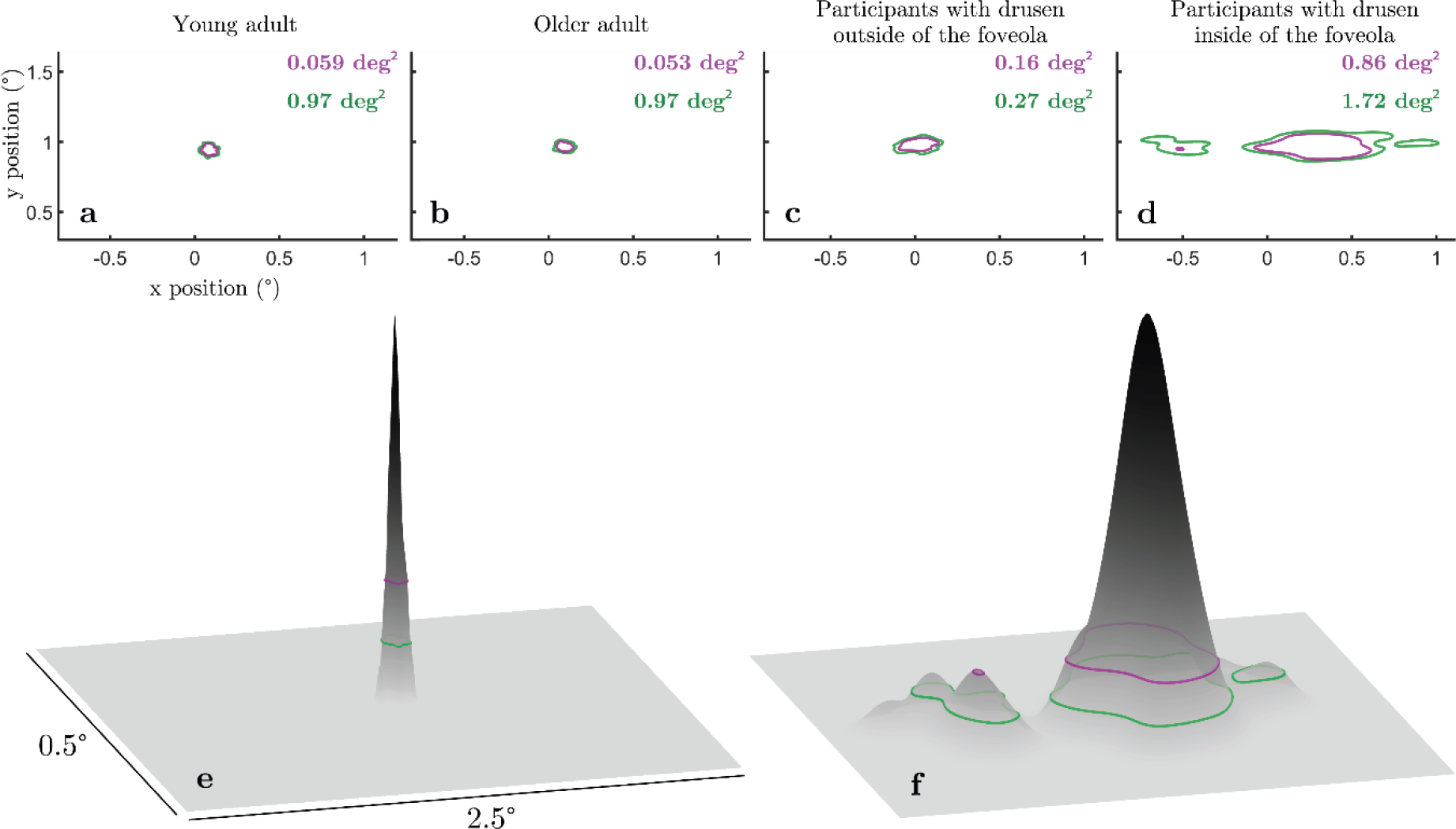
Impact of drusen on fixation stability. (a) The top row shows sample data respectively from a young adult, an older adult, a participant with foveal drusen and a participant with foveolar drusen. **(b)** Probability density of the young adult participant of (a) on the left and the foveolar participant of (a) on the right. ISOA contours encompassing 68% (purple) and 86% (green) of the eye position data.

**Figure 7.**
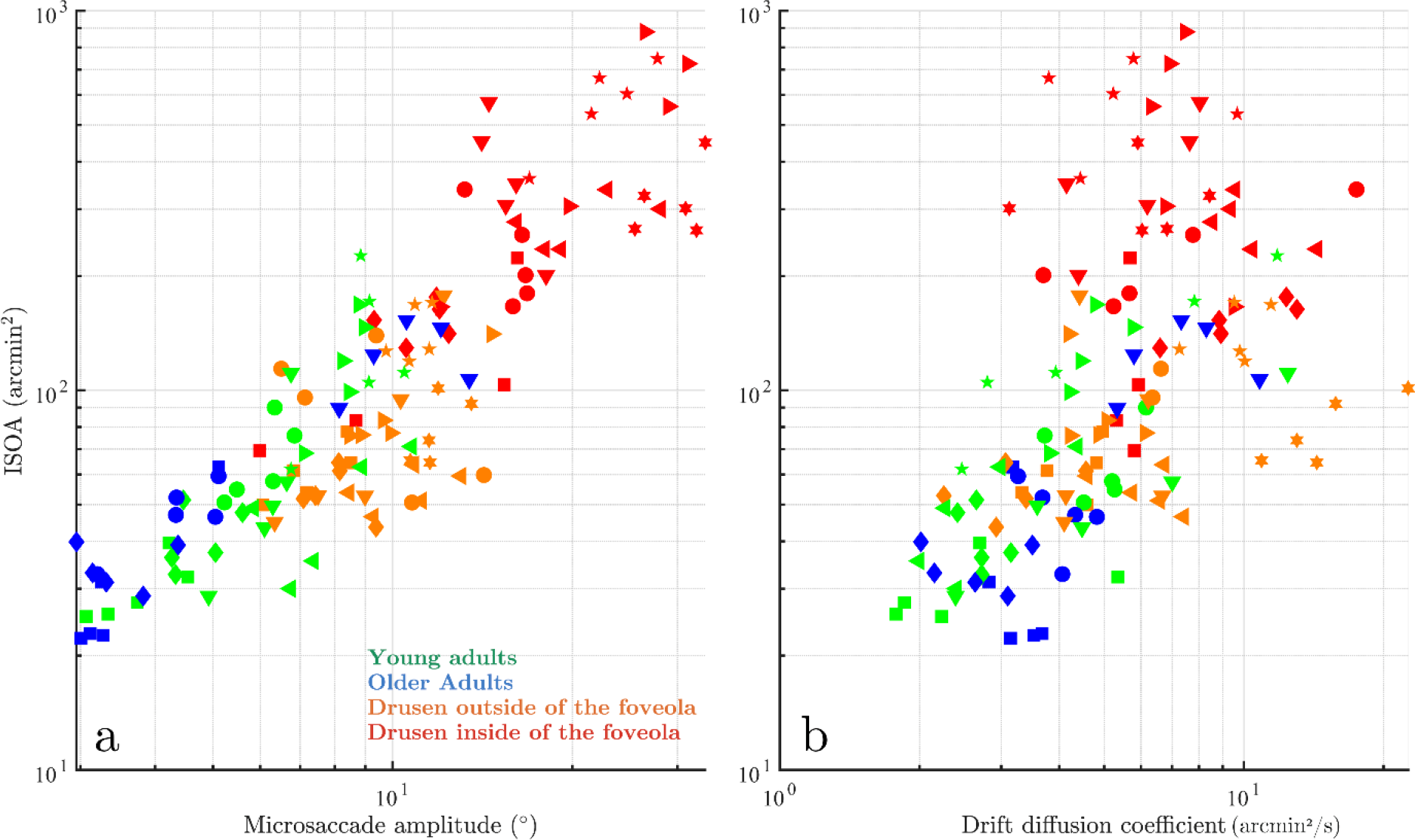
Contribution of microsaccade amplitude and drift coefficient to fixation stability. (a) Microsaccade amplitude vs. 86% ISOA. **(b)** Drift coefficient vs. ISOA on the right. One symbol represents one trial of a participant from its colored group.

## Discussion

The objective of the present study was to see whether structural retinal changes in presymptomatic AMD can have detectable consequences on functional aspects of vision. More specifically, we tested the hypothesis that FEM abnormalities are associated with early structural retinal anomalies, such as the appearance of small central drusen.

The primary finding of this study is that participants with foveal drusen exhibit significantly larger microsaccades and higher fixation instability (measured by ISOA) as compared to healthy controls, while maintaining equivalent microsaccade rates. Strikingly, drusen eccentricity with respect to the center of the fovea appears to be the main determinant factor associated with larger microsaccades and ISOAs. Indeed, we found that the more central the drusen were, the more unstable the fixation was; conversely, the further from the center of the fovea the drusen were, the more the participants tended to behave like controls, with minimal to no changes in fixation stability. We propose the following quantification criterion to assess the point after which drusen eccentricity strongly affect fixation characteristics: Microsaccade amplitude starts increasing exponentially when average drusen eccentricity is less than 1° with respect to the center of the fovea. Considering the size of the typical drusen and of the fixation target, the threshold point approximately corresponds to the size of the foveola, i.e. the 350 µm foveal pit as defined by Kolb^40^. Our results further suggest that quantification of microsaccade amplitude and ISOA in a simple ocular fixation test could in principle be sufficient to discriminate healthy participants from those with retinal anomalies in the foveola, such as drusen. Previous studies in patients with central scotomas suggest that these patients may develop one or more eccentric areas of fixation (referred to as preferred retinal locus or PRL)^41^, giving rise to an extended overall fixation area. We therefore checked in our presymptomatic subjects from the DIF group whether they express increased and possibly multipeaked fixation densities (as opposed to Gaussian-like distributions expected for the other groups). This was indeed the case, as only in the DIF group the ISOA contours exhibited multiple islands of fixation (see Fig. 6). In this study, we also evaluated the impact of foveal drusen density and eccentricity on the ocular diffusion drift coefficient, which provides a displacement estimator reflecting the randomness and stability of visual fixation^35^. The motion spectrum of drift epochs without microsaccades appeared however impacted by the presence of drusen and their centrality, especially in the middle frequency range. The dynamics of the complete eye movements (i.e. microsaccades included) during fixation appears successively altered with age, with the presence of foveal and further with foveolar drusen. This confirms the impact of drusen on microsaccade dynamics, and the importance to finely characterize them. Further research is needed to understand the role and implication of drusen on visual sampling and perception.

Another interesting observation stemming from this study concerns the definition of *healthy older adults* from an ophthalmological point of view. Twenty participants of this study were initially recruited as healthy controls based on the fact that (i) no visible foveal drusen were found using standard SD-OCT imaging analyzed by ophthalmologists, and (ii) no visual symptoms (in terms of visual acuity, contrast sensitivity, and so on) were observed during inclusion to the Silversight cohort. Subsequently, using AO-FIO-based gaze-dependent retina imaging, 45% of these participants were found to actually have small drusen in their fovea. This experimental observation, made with the use of more advanced imaging technique, aligns with early studies suggesting that drusen are more prevalent than previously assumed^42,43^. It also emphasizes the utmost importance of the development of sensitive measurement techniques that can be applied at presymptomatic stages to detect the presence of small drusen. While recent progress in retinal imaging resulted in retromode^44^ or gaze-dependent^16^ methods, which can both detect drusen invisible in standard SD-OCT images, this study opens up a new avenue in drusen detection research by showing that the appearance of small drusen can be detected based on simple functional measures, i.e. by looking at the FEM. Moreover, the discovered link between central drusen and fixation stability can explain the discrepancy present in the literature regarding the differences in fixation stability between young and healthy older adults, with some studies reporting an age-related decrease in stability^45–47^ and other studies, including ours, reporting no effect of age^48–50^. If the presence of foveal drusen is a common trait among older participants and these drusen do affect fixation stability, as we observe in our experiment, then the putative effect of age on fixation stability in the above studies can in fact be explained by the presence of participants with undetected central drusen in the sample of older adults (since virtually all previous studies used standard retina fundus imaging or SD-OCT). In conclusion, a combination of imaging methods and functional oculomotor measures, with a potential addition of subjective patient-reported measures of retinal health^50^ can be part of the general strategy for multifactor screening for retinal diseases, including AMD, in the general population. DIF participants were clearly clustered together around higher values of both ISOA and microsaccade amplitudes, suggesting the possibility of automatic detection procedure for the existence of foveolar drusen that could for instance use both ISOA and microsaccade amplitude values. While it is currently impossible to know whether or not existing drusen will develop into severe forms of AMD^51^, detection of drusen at the earliest possible stage will allow to monitor their progression and take the necessary precautions (e.g. apply anti-VEGF treatment^4^) before critical and irreversible damage occurs. Health style changes might help slow down their growth, and new drugs are currently being clinically tested to prevent the formation and evolution of drusen^7^.

To the best of our knowledge, alterations of fine oculomotor control in relation to small, asymptomatic changes of retinal structure have never been previously reported, whereas the assessment of eye movements to identify non-invasive biomarkers of a number of diseases is not new^52^. Research on eye movements has indeed existed for more than a century, and infrared video eye trackers capabilities, accuracy, and accessibility have renewed the interest on eye movements to clinicians and psychologists. FEM have been explored in neurological disorders such as amblyopia^53,54^, schizophrenia^55^, Parkinson’s disease^56^, and others^57^. There have also been attempts to reveal new FEM-based biomarkers to predict multiple sclerosis and follow its progression^58^. However, when it comes to detecting subtle retinal structural anomalies by means of FEM analysis, it is key to be able to record the whole spectrum of microsaccade amplitudes, down to 1 arcminute. The recent use of adaptive optics retinal imaging instruments has made it possible to reach this resolution. The AO-FIO used in this study allowed us to observe an average healthy microsaccade amplitude of 6.3 arcminutes, whereas participants with drusen had an average microsaccade amplitude of 13.8 arcminutes. In the past 20 years, FEM studies using pupil-based eye trackers have defined a microsaccade with amplitudes up to 1-2°^25,59–61^, and basic fixation tasks often showed average microsaccade amplitudes of 20-60 arcminutes. It is thus possible that the differences between controls and drusen participants found in our study would not have been detected using these less precise eye trackers and/or a fixation target larger than ours, because microsaccade amplitude is known to be target-dependent^23^. Further studies are being conducted in our team to confirm such hypotheses. This study, like many in the past decade, shows that FEM reveal a lot of unique information about patients. The system we use is probably too advanced and complex for immediate clinical use but the study is paving the road towards future FEM analysis in ophthalmologic testing with future accessible tracking methods that are starting to become available in the market.

While general functional significance of microsaccades is still a matter of debate, many researchers agree that microsaccades play an important functional role in the presence of foveolar stimuli, and the foveola size is even considered as a reference to define microsaccades^59,62^. As discussed in Poletti’s review on attention and eye movements at the foveal scale^59^, any stimuli in the foveola can be attention-grabbing and whether it is voluntary or not, it is possible to have microsaccade directed towards different stimuli as close as 20 arcminutes from each other. It would not be correct to consider foveal drusen as inputs to the visual system, but our data suggest that they might change retinal processing in specific retinal locations. Drusen appears to distort foveal vision but the exact role of or consequence on the oculomotor system remains to be determined. Fixation instability could be the result of the oculomotor system adaptation to avoid drusen locally. The second hypothesis consists in a more widespread RPE/photoreceptor dysfunction and degraded foveal sensory integration which would in turn impair motor control. The local compensatory versus global oculomotor deficiency hypotheses require further investigation. The relative impact of each eye’s drusen on binocular vision is another open subject of research to supplement these findings.

In conclusion, by combining state-of-the-art retinal imaging techniques and novel retinal motion tracking approach, we report an association between the presence of foveal drusen and fixation instability, hence linking structural and functional changes during aging. By demonstrating a strong association between foveal drusen properties and oculomotor behavior in a simple ocular fixation task our results suggest that FEM can be considered as a functional biomarker for innovative AMD screening approaches. More generally, our results suggest a possibility that the presence of even small drusen may affect visual perception. Thus, further studies are required to test potential effects of drusen on performance in tasks requiring high visual acuity, contrast sensitivity and photoreceptor function.

## Supporting information

Clip 1

Clip 2

## Abbreviations

FEM: fixational eye movements
AMD: age-related macular degeneration
RPE: retinal pigment epithelium
AO-FIO: adaptive optics flood illumination ophthalmoscope
RMS-S2S: root mean square sample-to-sample
ISOA: isoline area
SD-OCT: spectral domain optical coherence tomography

