## Supplementary figures and images for "Foveolar drusen decrease fixation stability in pre-symptomatic AMD"

### Sup 1

#### Power spectra accross groups - vertical motion

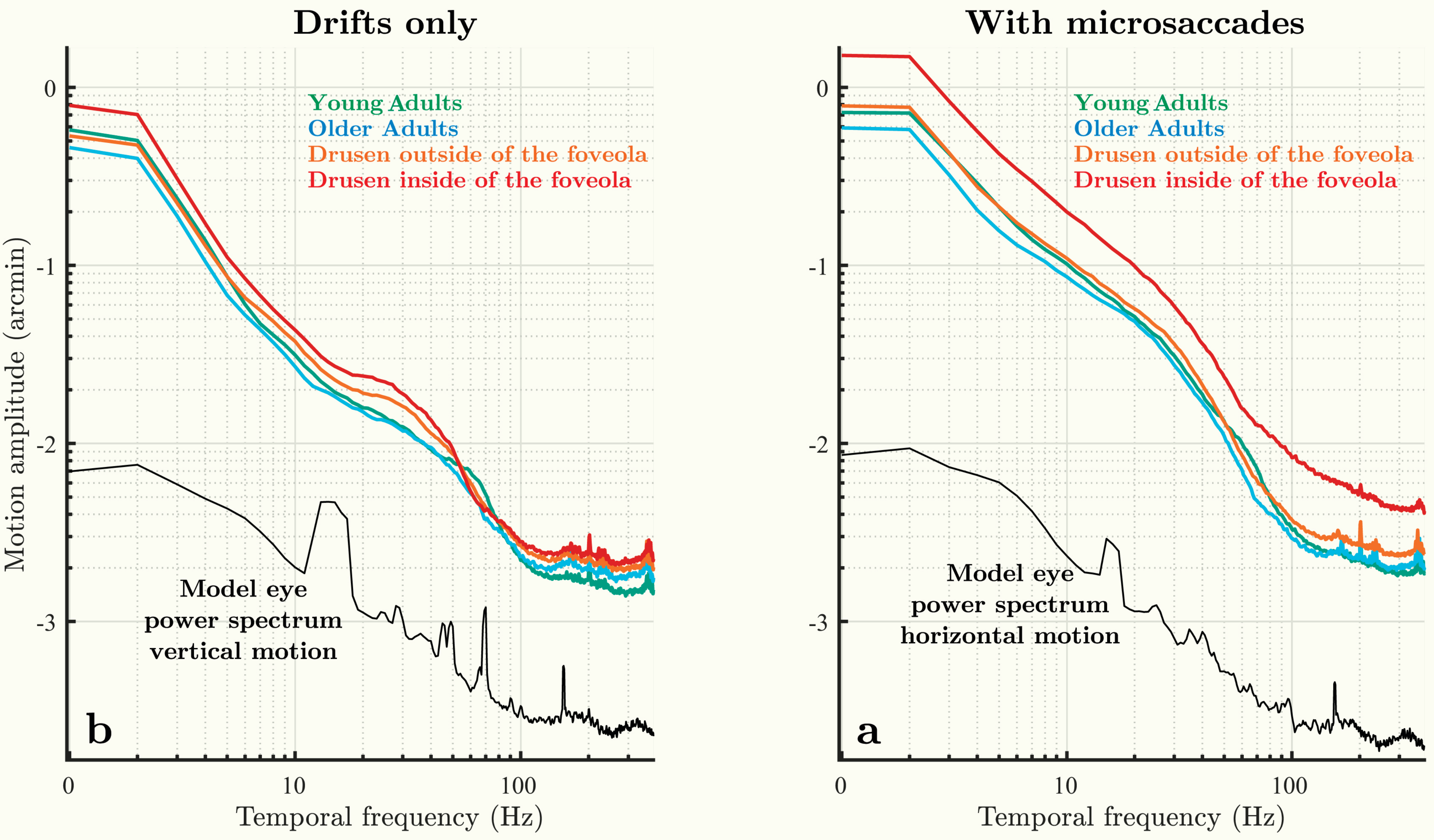

### Sup 2

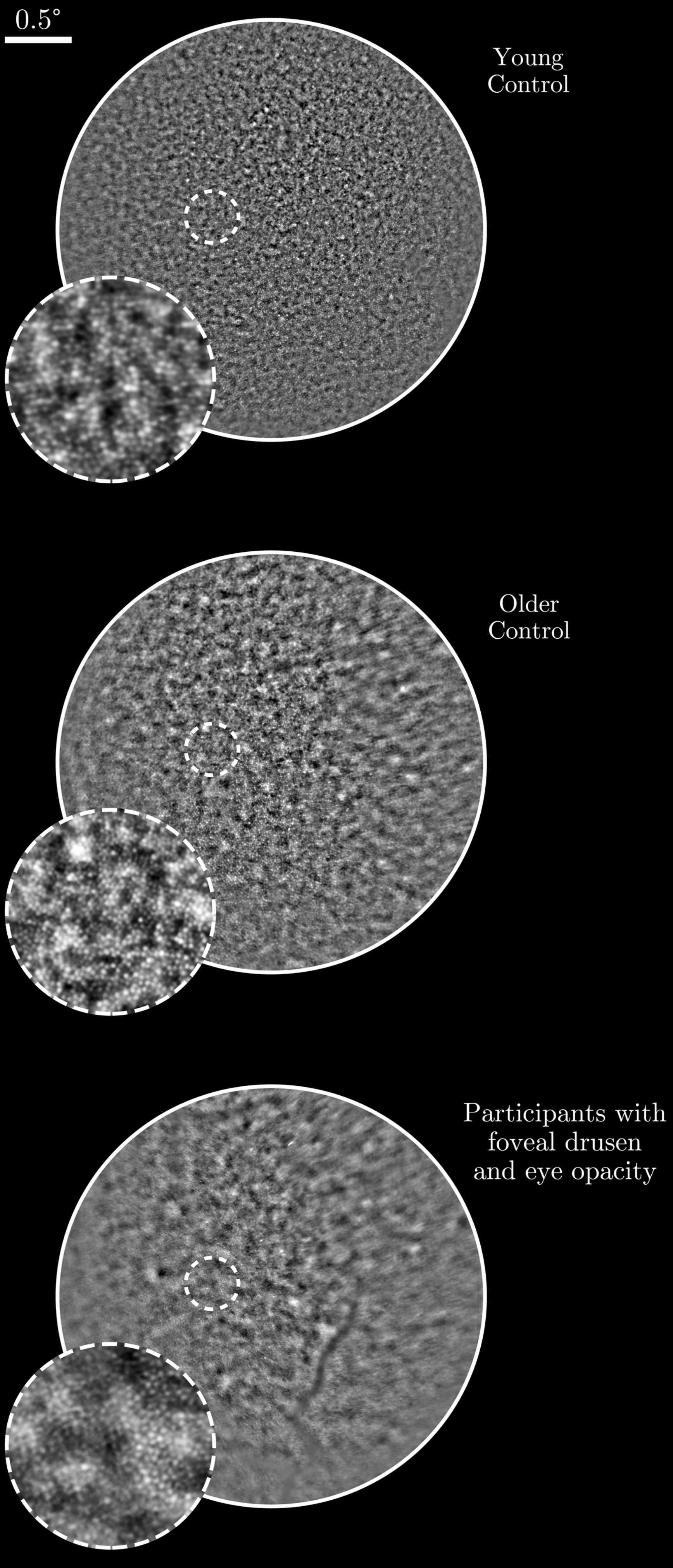
